# Osteoblast-osteoclast co-culture amplifies inhibitory effects of FG-4592 on osteoclast formation and reduces bone resorption activity

**DOI:** 10.1101/863498

**Authors:** Philippa A Hulley, Ioanna Papadimitriou-Olivgeri, Helen J Knowles

## Abstract

The link between bone and blood vessels is regulated by hypoxia and the hypoxia-inducible transcription factor, HIF, which drives both osteogenesis and angiogenesis. The recent clinical approval of PHD enzyme inhibitors, which stabilise HIF protein, introduces the potential for a new clinical strategy to treat osteolytic conditions such as osteoporosis, osteonecrosis and skeletal fracture and non-union. However, bone-resorbing osteoclasts also play a central role in bone remodelling and pathological osteolysis and HIF promotes osteoclast activation and bone loss in vitro. It is therefore likely that the final outcome of PHD enzyme inhibition in vivo would be mediated by a balance between increased bone formation and increased bone resorption. It is essential that we improve our understanding of the effects of HIF on osteoclast formation and function, and consider the potential contribution of inhibitory interactions with other musculoskeletal cells.

The PHD enzyme inhibitor FG-4592 stabilised HIF protein and stimulated osteoclast-mediated bone resorption, but inhibited differentiation of human CD14+ monocytes into osteoclasts. Formation of osteoclasts in a more physiologically relevant 3D collagen gel did not affect the sensitivity of osteoclastogenesis to FG-4592, but increased sensitivity to reduced concentrations of RANKL. Co-culture with osteoblasts amplified inhibition of osteoclastogenesis by FG-4592, whether the osteoblasts were proliferating, differentiating or in the presence of exogenous M-CSF and RANKL. Osteoblast co-culture dampened the ability of high concentrations of FG-4592 to increase bone resorption.

This data provides support for the therapeutic use of PHD enzyme inhibitors to improve bone formation and/or reduce bone loss for treatment of osteolytic pathologies, and indicates that FG-4592 might also act to inhibit the formation and activity of the osteoclasts that drive osteolysis.

## Introduction

The skeletal system and the vasculature are strongly linked during bone development, skeletal aging and in bone pathologies such as osteoporosis, osteonecrosis and skeletal fracture and non-union [1]. Skeletal cells secrete angiogenic factors to stimulate new blood vessel formation. In turn skeletal blood vessels supply osteoblastic bone-forming cells with oxygen, nutrients, growth factors and essential mineralisation components such as calcium and phosphate, maintain stem and progenitor cells and regulate skeletal cell behaviour.

The link between blood vessels and bone is regulated by hypoxia and the hypoxia-inducible transcription factor, HIF [2, 3]. HIF comprises an inducible alpha subunit (HIF-1α, HIF-2α) and a constitutively expressed beta subunit. Under standard conditions HIF-α is post-translationally hydroxylated by the prolyl-4-hydroxylase enzymes (PHD1–3), targeting it for interaction with the von Hippel–Lindau (VHL) protein and proteasomal degradation. The PHD enzymes are governed by O_2_ availability and so exhibit reduced activity under hypoxia. HIF-α then accumulates, translocates to the nucleus, dimerizes with HIF-β and binds the hypoxia-response element to induce transcription of HIF target genes such as pro-angiogenic vascular endothelial growth factor (VEGF) (reviewed in [4]).

Genetic studies initially defined the HIF-driven link between osteogenesis and angiogenesis. Mice with osteoblast-specific overexpression of HIFα due to deletion of *Vhl* [2] or the *Phd1-3* enzymes [5, 6] over-express VEGF and develop dense, heavily vascularised long bones, whereas osteoblast-specific deletion of *Hif1a* or *Hif2a* produces the reverse phenotype [2, 7]. Similarly HIF stabilisation with PHD enzyme inhibitors increases vascularity and stimulates new bone formation, improving bone mineral density and bone strength in murine models of bone fracture [8–11], distraction osteogenesis [12] and osteoporosis [13, 14]. The recent approval of novel and specific PHD enzyme inhibitors for clinical use [15, 16] therefore introduces the potential for a new strategy to treat osteolytic diseases.

However, many questions still need to be answered. For instance, some effects of the PHD / HIF pathway on bone are driven not by angiogenesis but by effects on bone-resorbing osteoclasts, which play a central role in bone remodelling and pathological osteolysis. Osteoclasts form by fusion of CD14+ monocytic precursors, induced by the cytokines macrophage colony-stimulating factor (M-CSF) and receptor activator of nuclear factor kappa B ligand (RANKL), to produce multi-nucleated bone-resorbing cells [17, 18]. By directly comparing HIF knockdown, HIF induction and PHD enzyme depletion in *in vitro* cultured murine and human osteoclasts we showed a striking role for HIF-1α and PHD2 in driving bone resorption by mature osteoclasts [19–22]. Osteoclast-specific inactivation of HIF-1α antagonises osteoporotic bone loss in mice, suggesting that HIF-1α also promotes osteoclast activation and bone loss *in vivo* [23]. Similarly, conditional deletion of *Phd2* in the monocyte/macrophage lineage causes reduced bone mass due to HIF-mediated production of erythropoietin, which inhibits osteoblast mineralisation and induces osteoclastogenesis and bone erosion [24].

However, effects of the PHD / HIF pathway on osteoblasts oppose its direct effect on osteoclasts. Mice with an osteoblast-specific mutation in *Phd2/3* display high bone mass without associated changes in vascularity, instead showing increased mRNA expression and elevated serum concentrations of osteoprotegerin (OPG), an inhibitor of osteoclast formation and activity [5].

Reduced numbers of osteoclasts are present in osteoblast-specific *Phd2/3^−/−^* mice *in vivo* and fewer osteoclasts form *in vitro* as a result of co-culture with *Phd2/3^−/−^* osteoblasts. This is driven predominantly by direct transcriptional effects of HIF-2α on *Opg* expression, which does not affect osteoblast proliferation or mineralisation [5]. We have observed elevated serum concentrations of OPG in *Phd3^−/−^* mice associated with reduced serum CTXI, indicative of reduced osteoclast activity *in vivo* [19].

It is likely that the final outcome of PHD enzyme inhibition *in vivo* would be mediated by a balance between increased bone formation and increased bone resorption. Increased trabecular bone mass in mice with an osteoblast-specific mutation in *Phd1-3* was described as the result of enhanced osteoclast-mediated bone resorption exceeded by increased bone formation [6]. This dampening of HIF-mediated osteoclast activation could encompass a moderate effect of HIF to delay cell fusion during osteoclast differentiation [19].

In light of the potential of HIF pathway activation as a therapeutic strategy to improve bone formation and/or reduce bone loss, it is essential that we also improve our understanding of the varied effects of HIF on osteoclast formation and function *in vivo*, especially considering the contribution of inhibitory interactions with other musculoskeletal cell types. This manuscript describes how 3-dimensional human osteoclast culture or osteoclast-osteoblast co-culture can be used to model *in vitro* some of the non-angiogenic effects of PHD enzyme inhibition *in vivo* and how it could facilitate detailed evaluation of the molecular mechanisms involved.

## Results

### FG-4592 stimulates osteoclast bone resorption but reduces differentiation

Of the classical non-specific PHD enzyme inhibitors which include dimethyl oxalyl glycine (DMOG), L-mimosine, desferrioxamine and CoCl_2_, DMOG causes the greatest increase in osteoclast-mediated bone resorption, despite some evidence of toxic effects [19, 22]. DMOG is commonly used *in vitro* at ≤1 mM concentration to stabilise HIF and activate HIF-regulated transcription. FG-4592, a more specific PHD enzyme inhibitor that recently obtained clinical approval [16], was titrated to stabilise equivalent quantities of HIF protein in comparison with DMOG (Figure 1a). A 40-fold lower concentration of FG-4592 than DMOG induced equivalent HIF transcriptional activation and expression of HIF-regulated proteins such as Glut-1 (Figure 1a, b). ‘HIF-equivalent’ doses of FG-4592 induced an increase in bone resorption comparable to DMOG (Figure 1c, d, f), although without any reduction in osteoclast number (Figure 1e, f).

**Figure 1:**
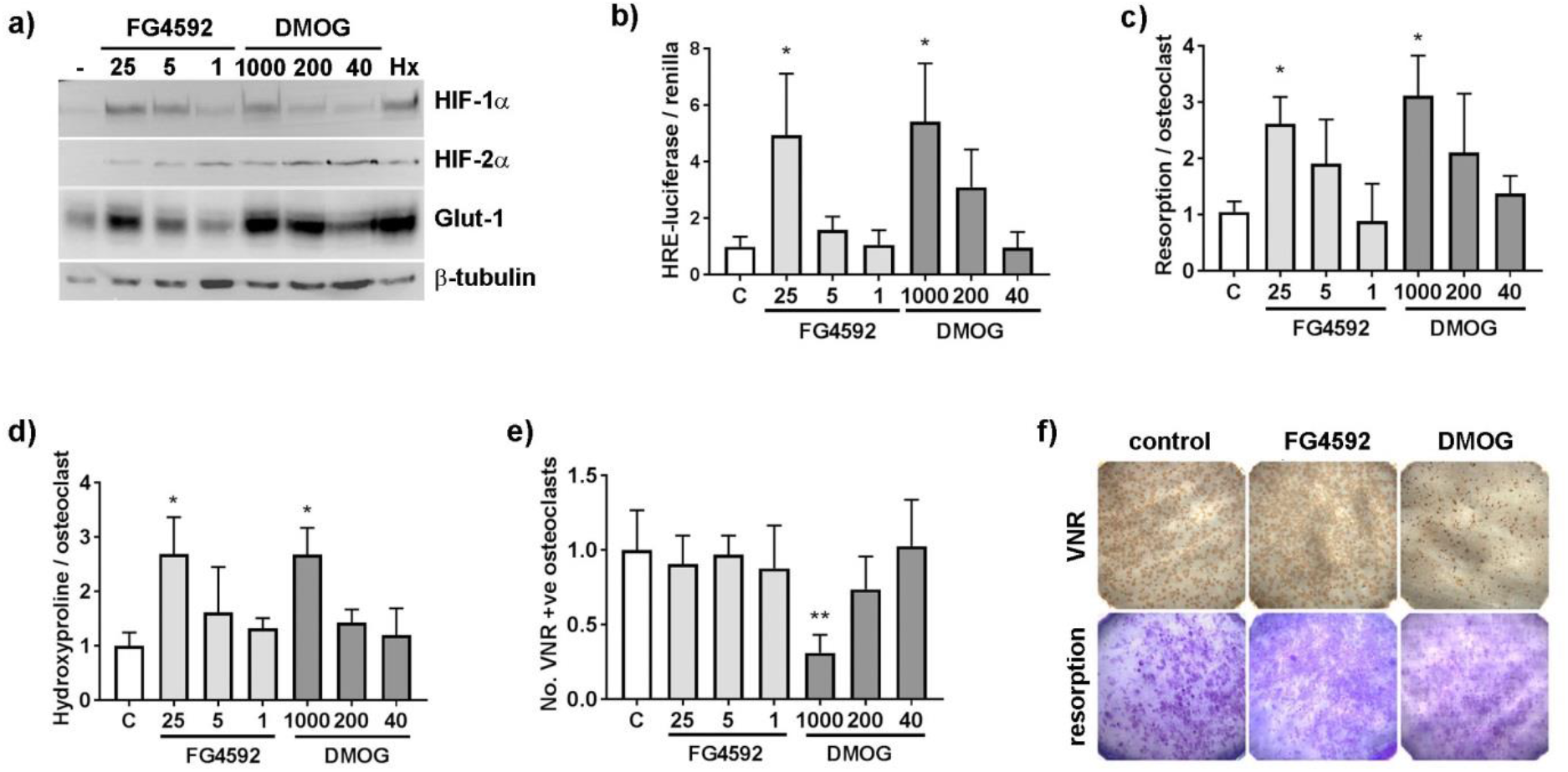
FG-4592 stimulates osteoclast bone resorption.

(a) Western blot showing stabilisation of HIF-1α and HIF-2α protein and induction of Glut-1 and (b) HRE-luciferase activity (n=5) in response to 24 h exposure of mature osteoclasts to FG-4592 (1-25 μM), DMOG (40-1000 μM) or hypoxia (2% O_2_). (c-e) Quantified effect of 24 h exposure of mature human osteoclasts cultured on dentine to FG-4592 and DMOG with respect to (c) the area of dentine resorbed per osteoclast (n=4), (d) the amount of hydroxyproline released per osteoclast (n=4) and (e) osteoclast survival (number of VNR-positive osteoclasts present; n=5). (f) Representative images of dentine discs after VNR immunohistochemistry or toluidine blue staining of resorption tracks. *, p<0.05; **, p<0.01.

Effects of FG-4592 were next analysed on osteoclast differentiation. FG-4592 reduced the rate of proliferation of CD14+ monocyte precursors in the presence of M-CSF (Figure 2a). FG-4592 also reduced osteoclast formation on both plastic (Figure 2b, c) and dentine (Figure 2d) to 23.3-57.9% (25 μM) and 58.1-84.1% (5 μM) of control, potentially due to reduced proliferation of the precursor population. Interestingly, those osteoclasts that did form in the presence of 25 μM FG-4592 exhibited a greatly increased capacity for bone resorption (Figure 2e). DMOG caused much greater inhibition of both monocyte proliferation and osteoclast formation, likely due to cumulative toxicity over the 9 day experimental period, and was therefore not used for further experiments.

**Figure 2:**
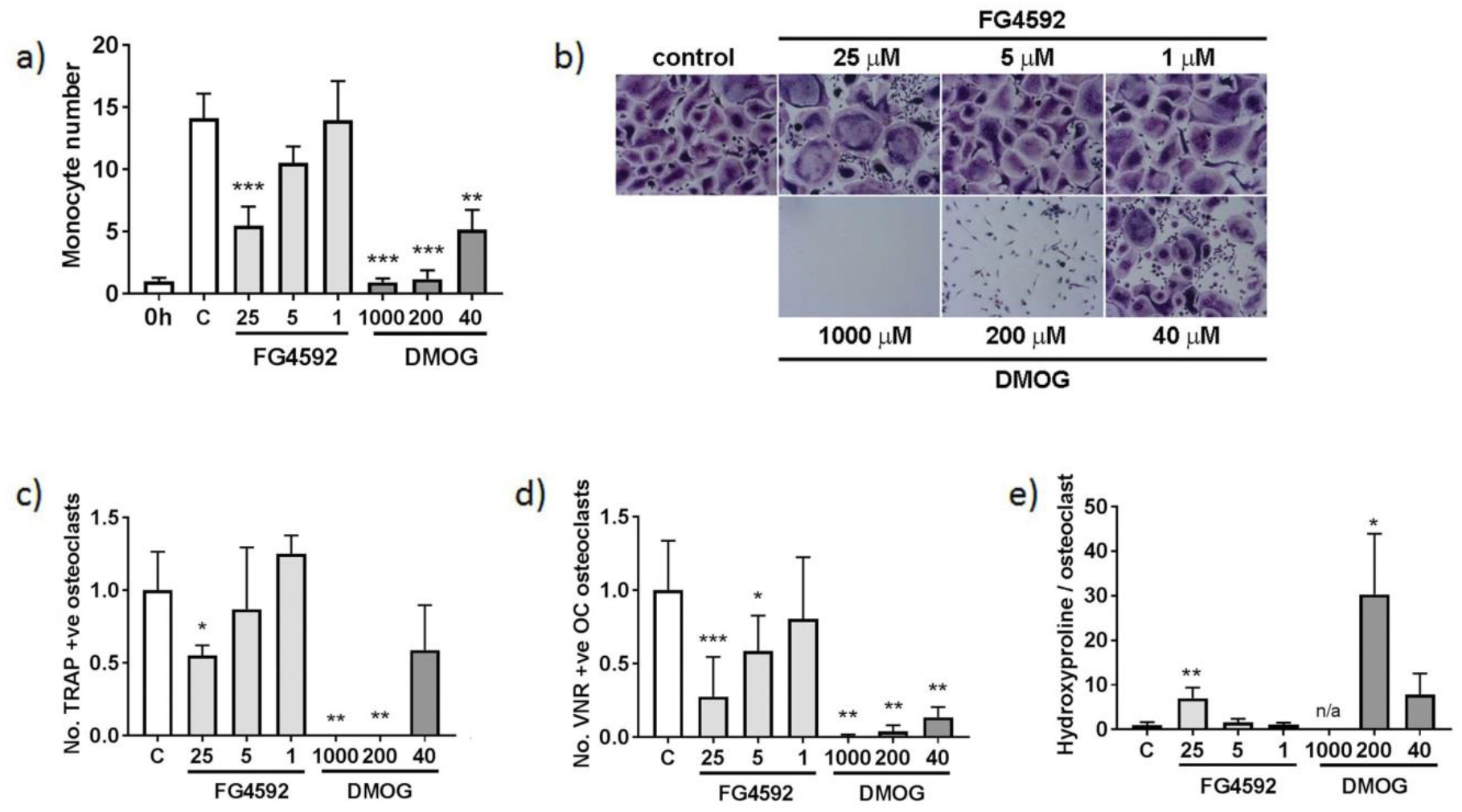
FG-4592 inhibits osteoclast differentiation.

(a) Relative number of CD14+ monocytes after 9 days exposure to FG-4592 (1-25 μM) or DMOG (40-1000 μM) (n=6). Significance in comparison to the M-CSF (C) control. (-) = untreated control. (b) Representative images of TRAP staining of mature osteoclasts on cell culture plastic. (c-e) Quantified effect of 9 days exposure of mature human osteoclasts to FG-4592 and DMOG with respect to (c) the number of multi-nucleated TRAP-positive osteoclasts formed (n=4), (d) the number of VNR-positive osteoclasts formed on dentine (n=6) and (e) the amount of hydroxyproline released from dentine per osteoclast from day 7 to day 9 (n=6). *, p<0.05; **, p<0.01; ***, p<0.001.

### 3D culture increases sensitivity of osteoclastogenesis to RANKL but not to FG-4592

We next investigated whether differentiating osteoclasts in a more physiologically relevant 3D culture system might alter effects of PHD enzyme inhibition. By light microscopy, osteoclasts differentiated inside a 2 mg/ml collagen gel exhibited an equivalent rate of differentiation to those cultured on plastic (Figure 3a). Immunohistochemical analysis of the osteoclasts formed within the gel revealed them to be multi-nucleated and express the osteoclast markers TRAP and cathepsin K (Figure 3b). These osteoclasts were able to form resorption tracks after release from the gel and re-seeding onto dentine discs (Figure 3b). Following gel release, mature osteoclasts were re-seeded onto cell culture plastic for quantification of the number of TRAP-positive osteoclasts formed during 3D differentiation in the presence of FG-4592 (Figure 3c). No change in sensitivity of osteoclast formation to FG-4592 was observed as a result of 3D culture (Figure 3d). We did however observe an increased sensitivity of osteoclastogenesis to reduced RANKL concentrations in 3D. A 54.7% reduction in osteoclast formation was evident in 3D versus 2D culture at 5 ng/ml RANKL (Figure 3e, p<0.01). It was not possible to determine direct effects of either treatment on osteoclast activity, as the 6 day post-reseeding period necessary to visualise resorption tracks obscured effects of 3D culture.

**Figure 3:**
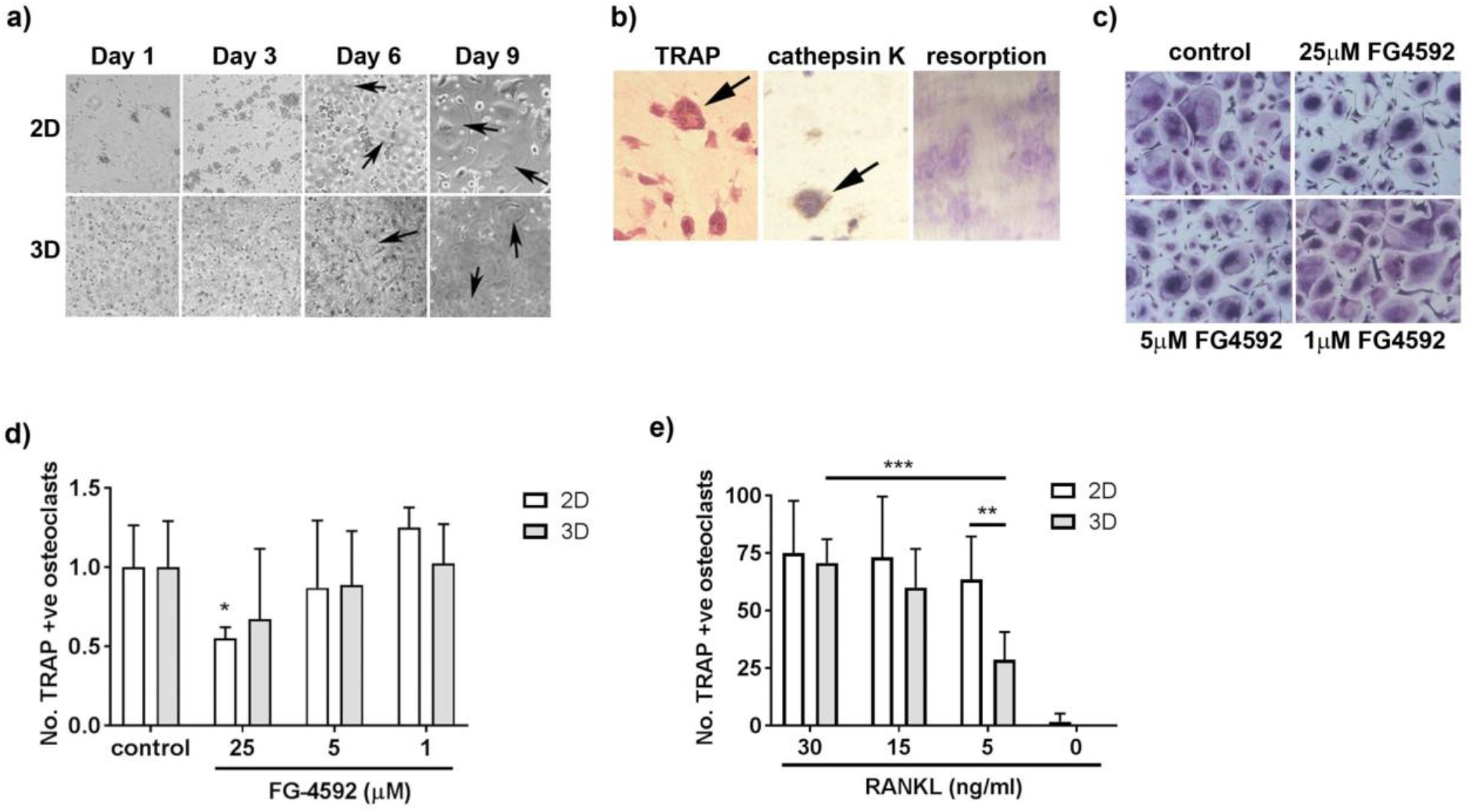
3D culture alters effects of RANKL on osteoclastogenesis.

(a) Representative brightfield image of osteoclast differentiation in 2D versus 3D culture. (b) Staining of formalin-fixed paraffin-embedded 3D gel for TRAP (left panel) and cathepsin K (middle panel). Arrows indicate multi-nucleated osteoclasts. Right panel: resorption tracks formed by re-seeded osteoclasts on dentine discs. (c) TRAP staining of multi-nucleated osteoclasts differentiated in collagen gels in the presence of FG-4592 then re-seeded in 2D culture. (d) Quantification of osteoclasts differentiated in collagen gels with FG-4592 (n=6) or (e) varied concentrations of RANKL (n=4) then re-seeded in 2D culture. Comparison with osteoclasts differentiated in 2D culture. *, p<0.05; **, p<0.01; ***, p<0.001

### Co-culture with osteoblasts alters effects of FG-4592 on osteoclastogenesis

We next considered indirect effects of FG-4592 on osteoclast formation when monocytes were co-cultured with osteoblasts. To enable comparison with the direct effects of FG-4592 on osteoclastogenesis seen in Figure 2, we first cultured primary human osteoblasts with CD14+ monocytes for 9 days in the presence of exogenous M-CSF and RANKL. Multi-nucleated TRAP-positive osteoclasts formed in the co-culture system (Figure 4a) which displayed greater sensitivity to lower doses of FG-4592 (Figure 4b). Due to difficulties visualising TRAP-positive osteoclasts in some co-cultures, lysates were made from osteoclasts or co-cultured cells and analysed for cathepsin K activity, an osteoclast marker enzyme that readily distinguishes mature cells from their monocytic precursors (Figure 4c). A marked increase in inhibition of osteoclast formation by FG-4592 was again observed (Figure 4d). This was not due to over-population of the culture with osteoblasts, which showed a reduced rate of proliferation in the presence of FG-4592 (Figure 4e). Similarly, the osteoclasts that formed in the presence of 25μM FG-4592 showed dampened resorption activity when co-cultured with osteoblasts (Figure 4f).

**Figure 4:**
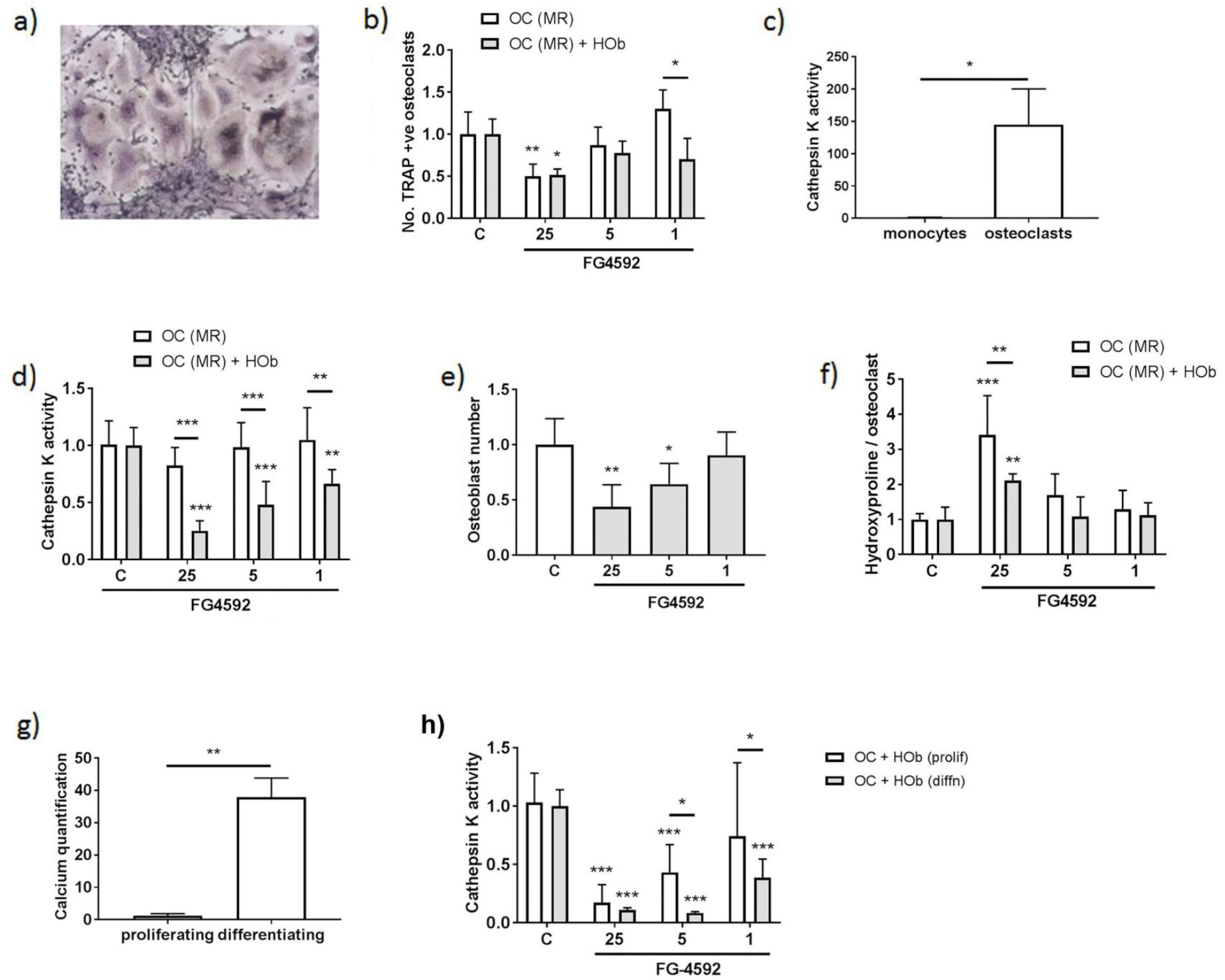
Monocyte-osteoblast co-culture amplifies inhibition of osteoclastogenesis by FG-4592.

(a) Representative brightfield image of osteoclast differentiation following monocyte-osteoclast co-culture in the presence of M-CSF and RANKL. (b) Relative effect of FG-4592 (1-25 μM) on the number of multi-nucleated TRAP-positive osteoclasts formed in osteoclast monoculture versus monocyte-osteoclast co-culture in the presence of M-CSF and RANKL (n=4). (c) Relative cathepsin K activity measured in monocyte and osteoclast lysates (n=4). (d) Relative effect of FG-4592 on cathepsin K activity in osteoclast monoculture versus monocyte-osteoclast co-culture in the presence of M-CSF, RANKL (n=5). (e) Relative number of osteoblasts after 9 days exposure to FG-4592 (1-25 μM) (n=6). (f) Relative effect of FG-4592 on the amount of hydroxyproline released from dentine (normalised to cathepsin K activity as a reflection of osteoclast numbers) from day 7 to day 9 by osteoclast monocultures versus monocyte-osteoclast co-cultures in the presence of M-CSF and RANKL (n=6). (g) Relative calcium concentration in cell pellets from differentiated versus proliferating osteoblasts as a measure of mineralisation (n=3). (h) Relative effect of FG-4592 on cathepsin K activity in monocyte-osteoclast co-cultures in the absence of M-CSF, proliferating versus differentiating (50 μg/ml ascorbate, 5 mM β-glycerophosphate) osteoblast cultures (n=6). *, p<0.05; **, p<0.01; ***, p<0.001.

M-CSF and RANKL, as well as other pro-and anti-osteoclastogenic factors, are supplied *in vivo* by cells within the bone microenvironment such as osteoblasts, osteocytes and T cells. Co-culture with osteoblasts is therefore able to stimulate osteoclastogenesis over a 14-21 day period in the absence of exogenous M-CSF or RANKL [25, 26]. Treatment with FG-4592 had a similar magnitude of effect on osteoclastogenesis during extended co-culture with proliferating primary human osteoblasts alone as with the shorter co-culture period in the presence of exogenous cytokines (cathepsin K activity, Figure 4g; compare with Figure 4c). The observed level of inhibition with FG-4592 was magnified by co-culture with differentiating, rather than proliferating, osteoblasts (Figure 4g,h).

## Discussion

PHD inhibitors have now entered clinical use, increasing the feasibility of using HIF pathway activation as a therapeutic strategy to improve bone formation and/or reduce bone loss. However, the complexity of their effects on osteogenic-angiogenic coupling and the potential consequences of actions on the HIF pathway affecting osteoclast:osteoblast interactions are far from clear. This manuscript describes how the selective PHD enzyme inhibitor FG-4592 reduces osteoclast differentiation but enhances osteoclast-mediated bone resorption *in vitro* in 2D monoculture. While 3D monoculture did not alter the effect of FG-4592 on osteoclast differentiation, 2D co-culture with osteoblasts caused marked inhibition of osteoclastogenesis by FG-4592 as well as diminished stimulation of bone resorption.

It is of interest that FG-4592 caused a comparable increase in osteoclast-mediated bone resorption to DMOG under standard monoculture conditions. DMOG is a 2-oxoglutarate analogue which acts as a broad spectrum inhibitor of 2-oxoglutarate-dependent dioxygenases, a family that includes the HIF-regulating PHD enzymes. DMOG is generally a better mimic of hypoxic transcriptional responses than selective PHD enzyme inhibitors as it includes transcriptional responses mediated by enzymes such as factor inhibiting HIF (FIH) [27]. This means that although we titrated the inhibitors to find concentrations that stabilised equivalent amounts of HIF, with comparable induction of Glut-1 protein and PGK1 HRE luciferase activity, the full panel of downstream target genes induced is likely to diverge. Other groups have observed that FG-4592 induces expression of HIF-regulated VEGF and Glut-1 *in vitro* over a 5-100 μM dose range [28–31], equivalent to the concentrations used in this study. FG-4592 also induces HIF-mediated expression of erythropoietin and inhibits expression of hepcidin as part of its function to treat anaemia in patients with chronic kidney disease [16]. The equivalent effect of FG-4592 and DMOG on osteoclast activity suggests that effects of hypoxia on osteoclast-mediated bone resorption are driven predominantly by the PHD enzymes, as we have previously reported [19, 22].

Although 3D collagen gel culture did not alter direct effects of FG-4592 on osteoclastogenesis, it did increase sensitivity to changes in RANKL concentration. We could find no reference in the literature regarding effects of 3D culture on osteoclast biology, although it is known to affect some differentiation and drug-responses in other cell types. We have previously described enhanced differentiation of primary rat osteoblasts and MC3T3-E1 cells in 3D versus 2D culture, as well as increased sensitivity to inhibition of proliferation by a protein kinase inhibitor [32]. 3D culture alters the sensitivity of prostate cancer cells to anti-neoplastic drugs [33] and enables spontaneous differentiation of mouse calvarial osteoblasts into osteocytes, in the absence of any other factors [34].

RANKL is a critical regulator of osteoclast differentiation and activity, whose expression can be regulated *in vivo* by HIF. HIF-1α increases RANKL expression in stromal fibroblasts [35], breast cancer cells [36] and osteocytic MLO-Y4 cells [37]. HIF-2α deficiency in mice enhances bone mass, in part by increasing osteoclastogenesis. HIF-2α stimulates RANKL-induced osteoclastogenesis via regulation of osteoclast Traf6 and also increases RANKL expression in osteoprogenitor cells [38]. This could mean that in 3D culture *in vivo*, possible effects of FG-4592 to drive a HIF-mediated increase in RANKL expression by other bone-resident cells could have large effects on osteoclast differentiation and activity within the local bone microenvironment.

As part of our 3D experiments, we developed a method enabling viable release and re-seeding of mature bone-resorbing osteoclasts from 3D collagen gel culture onto a new bone surface, which they proceeded to digest. Normally osteoclasts grown *in vitro* lose viability and bone-resorption activity when detached from the surface. Recent publications have achieved viable release of 2D-generated osteoclasts using accutase, although there is an indication that murine osteoclasts retain more viability than human osteoclasts following this procedure [39, 40]. The ability to viably move and re-seed mature bone-resorbing osteoclasts *in vitro* will enable osteoclast differentiation and activity to be studied separately experimentally, a separation which has proven impossible with current osteoclast culture techniques.

Co-culture of monocytes with osteoblasts is a well-known method of generating osteoclasts, utilising the RANKL supplied by the osteoblastic cells to induce differentiation. Our data that inhibition of osteoclast differentiation by FG-4592 was greater under co-culture with osteoblasts is similar to that previously described with reagents such as magnesium extract [41], triptolide [42] and Equisetum arvense [43]. It is possible that this is due to changes in the RANKL:OPG ratio, although our data regarding effects on OPG:RANKL expression was variable and inconclusive (data not shown). This could be because expression of both RANKL [35–38] and OPG [5, 19] can be induced by HIF activation and / or PHD enzyme inhibition. Full transcriptional investigation of this complex experimental system would facilitate identification of all contributing factors.

In summary, our data provides support for the use of PHD enzyme inhibitors as a potential therapeutic strategy to improve bone formation and/or reduce bone loss during treatment of osteolytic pathologies ranging from cancer and rheumatoid arthritis to osteoporosis and bone fracture. Our human primary cell co-culture data indicates that FG-4592 might act to inhibit the formation and activity of osteoclasts *in vivo*. This manuscript does not investigate whether it would, in parallel, stimulate angiogenesis and osteogenesis. However, FG-4592 inhibits the growth of melanoma and lung carcinoma tumours in murine models, in part by normalising tumour blood vessels [44], and increases vascularity in subcutaneous models of angiogenesis [28], partially via *in vivo* induction of VEGF [28, 30]. The recent approval of FG-4592 for clinical use [15, 16] therefore highlights the need to accelerate research into this potential new strategy to treat osteolytic disease.

## Methods

### Materials and ethical approvals

Reagents were obtained as follows: M-CSF (R&D Systems, Abingdon, UK), RANKL (Peprotech, London UK), DMOG (Cayman Chemicals, Michigan, USA), FG4592 (Selleckchem, Houston, USA). Unless stated other reagents were from Sigma-Aldrich (Gillingham, UK). Use of leucocyte cones for osteoclast differentiation and isolation of fibroblast-like synoviocytes from synovial tissue obtained from patients undergoing joint replacement surgery was approved by the London-Fulham Research Ethics Committee (11/H0711/7 and 07/H0706/81 respectively).

### Osteoclast culture

CD14+ monocytes were positively selected from the peripheral blood mononuclear cell component of leucocyte cones (NHS Blood and Transplant, UK) using magnetic CD14+ microbeads (Miltenyi Biotech, Surrey, UK). Monocytes were seeded onto dentine discs or plastic dishes in α-MEM (without ribonucleosides / deoxyribonucleosides) containing 10% FBS, 2 mM L-glutamine, 50 IU/ml penicillin and 50 μg/ml streptomycin sulphate. Osteoclastogenesis was induced by treatment with 25 ng/ml M-CSF and 30 ng/ml RANKL every 3–4 days for 9 days. Monocytes were maintained in 25 ng/ml M-CSF. Hypoxic exposure was at 2% O_2_, 5% CO_2_, balance N_2_ in a MiniGalaxy incubator (RS Biotech, Irvine, UK).

### 3D osteoclast culture

1×10^6^ CD14+ monocytes per well of a 24 well plate were pelleted, resuspended in 300 μl of 2 mg/ml collagen type I (Corning, New York, USA) and allowed to polymerise at 37°C. α-MEM and cytokines were applied as for standard osteoclast culture for 9 days. To release cells, gels were digested with 0.2 mg/ml collagenase type I at 37°C for 30 minutes, washed and resuspended in α-MEM prior to re-seeding onto plastic or dentine as required. Alternatively gels were formalin-fixed, paraffin-embedded and sectioned for analysis of internal cells by immunohistochemistry.

### Osteoclast assays

Tartrate-resistant acid phosphatase (TRAP) staining of formalin-fixed cells used naphthol AS-BI phosphate as a substrate, with reaction of the product with Fast Violet B salt. Multi-nucleated cells containing three or more nuclei were considered osteoclasts. Immunostaining for the vitronectin receptor (VNR) was with anti-CD51/61 (clone 23C6, 1:400; Bio-Rad, Oxford, UK) and for cathepsin K with a rabbit polyclonal antibody (3368-100, BioVision, California, USA). Measurement of cathepsin K activity as a surrogate for osteoclast number was performed in cells lysed in 1% Triton X-100 and assayed in an adaptation of previous methods [45, 46]. Briefly, 1 μg lysate was incubated with 100 μM Z-Gly-Pro-Arg-7-(4-methyl)-coumarylamide (MCA) (Bachem, St Helens, UK), a specific substrate for cathepsin K, in buffer [50 mM potassium phosphate buffer (pH 6.5), 2.5 mM DTT, 2.5 mM EDTA] for 1 h at 37°C. Generation of fluorescent product was assayed at 380 nm (excitation) and 450 nm (emission).

Resorption tracks on dentine discs were visualised by staining with 0.5% toluidine blue. Dentines were photographed, resorption tracks highlighted and the resorbed area quantified using ImageJ. Alternatively release of hydroxyproline was measured as an indication of bone resorption. Sample media was hydrolysed with HCl and dried, then hydroxyproline concentration was determined by reaction of chloramine-T oxidized hydroxyproline with 4-(dimethylamino)benzaldehyde (DAB) and quantification of the colorimetric product at 560nm.

### Co-culture with primary osteoblasts

Primary human osteoblasts were purchased from Sigma. Co-culture cells were titrated to determine the highest viable seeding density to allow continuous culture for the duration of the osteoclast differentiation assay. Co-culture cells were seeded at the determined density in 24 well plates and 1 × 10^6^ CD14+ monocytes were overlaid 24 h later. Osteoclastogenesis was induced either by treatment with 25 ng/ml M-CSF and 30 ng/ml RANKL every 3–4 days for 9 days, or with M-CSF and RANKL supplied by the osteoblasts over a 21 day period. Osteoblast differentiation (mineralisation) was induced over the same period by additional treatment with 50 μg/ml ascorbate and 5 mM β-glycerophosphate. Mineralisation was measured by calcium quantification [47]. Briefly, cells were lysed in 0.1% Triton X-100 then mineral was dissolved in 1.0 N acetic acid. Calcium levels were quantified with a working solution containing o-cresolphtalein complexone, with absorbance read at 570 nm.

### Western blots

Cells were homogenised in HIF lysis buffer (6.2 M urea, 10% glycerol, 5 mM dithiothreitol, 1% sodium dodecyl sulphate, protease inhibitors). Primary antibodies were against HIF-1α (clone 54, 1:1000; BD Biosciences, Oxford, UK), HIF-2α (ep190b, Novus Biologicals, Abingdon, UK), GLUT1 (ab14683, 1:2500; Abcam) or β-tubulin (clone TUB2.1, 1:2500).

### Luciferase assay

Osteoclasts were transfected with a PGK HRE–firefly luciferase plasmid (gifted by Professor AL Harris, Oxford, UK) and a pHRG–TK Renilla luciferase control plasmid (Promega, Southampton, UK) using Lipofectamine 2000 (Invitrogen). Luminescence was assayed after 24-28 h using the Dual-Luciferase Reporter Assay System (Promega), with firefly luciferase normalized to the Renilla transfection control.

### Metabolic assays

Glucose concentration in the media was measured using the Glucose (GO) Assay Kit. Mitochondrial dehydrogenase activity within the electron transport chain was assessed by adding Alamar blue (BioRad, Kidlington, UK) to cells in culture for 4 h. Results were normalized to osteoclast number. Relative cell number for mononuclear cells (non-osteoclasts) was determined using crystal violet staining.

### Realtime PCR

RNA was extracted in TRI reagent (Direct-Zol RNA Miniprep kit; Zymo Research, Irvine, CA, USA), reverse-transcribed and quantitative PCR was performed using Fast SYBR Green Master Mix in a Viia7 Real-Time PCR system (Applied Biosystems, Warrington, UK). Human primers were pre-validated Quantitect primers (Qiagen, Manchester, UK). Comparative quantification normalised target gene mRNA to β-actin (ACTB) mRNA.

### Statistical methods

Results are derived from at least three independent experiments. Data are presented as mean ± standard error and were analysed using GraphPad Prism. Statistical analysis comprised one-way or two-way ANOVA using Dunnett’s or Sidak’s multiple comparison as a post-hoc test. For experiments with only two conditions a t-test was applied. Results were considered significant at p < 0.05.

## References

1. Stegen S, Carmeliet G: The skeletal vascular system - Breathing life into bone tissue. Bone 2018, 115:50–58.

2. Wang Y, Wan C, Deng L, Liu X, Cao X, Gilbert SR, Bouxsein ML, Faugere MC, Guldberg RE, Gerstenfeld LC, Haase VH, Johnson RS, Schipani E, Clemens TL: The hypoxia-inducible factor alpha pathway couples angiogenesis to osteogenesis during skeletal development. J Clin Invest 2007, 117(6):1616–1626.

3. Yellowley CE, Genetos DC: Hypoxia Signaling in the Skeleton: Implications for Bone Health. Current osteoporosis reports 2019, 17(1):26–35.

4. Deng W, Feng X, Li X, Wang D, Sun L: Hypoxia-inducible factor 1 in autoimmune diseases. Cellular immunology 2016, 303:7–15.

5. Wu C, Rankin EB, Castellini L, Alcudia JF, LaGory EL, Andersen R, Rhodes SD, Wilson TL, Mohammad KS, Castillo AB, Guise TA, Schipani E, Giaccia AJ: Oxygen-sensing PHDs regulate bone homeostasis through the modulation of osteoprotegerin. Genes & development 2015, 29(8):817–831.

6. Zhu K, Song P, Lai Y, Liu C, Xiao G: Prolyl hydroxylase domain proteins regulate bone mass through their expression in osteoblasts. Gene 2016, 594(1):125–130.

7. Shomento SH, Wan C, Cao X, Faugere MC, Bouxsein ML, Clemens TL, Riddle RC: Hypoxia-inducible factors 1alpha and 2alpha exert both distinct and overlapping functions in long bone development. Journal of cellular biochemistry 2010, 109(1):196–204.

8. Shen X, Wan C, Ramaswamy G, Mavalli M, Wang Y, Duvall CL, Deng LF, Guldberg RE, Eberhart A, Clemens TL, Gilbert SR: Prolyl hydroxylase inhibitors increase neoangiogenesis and callus formation following femur fracture in mice. J Orthop Res 2009, 27(10):1298–1305.

9. Huang J, Liu L, Feng M, An S, Zhou M, Li Z, Qi J, Shen H: Effect of CoCl2 on fracture repair in a rat model of bone fracture. Molecular medicine reports 2015.

10. Stewart R, Goldstein J, Eberhardt A, Chu GT, Gilbert S: Increasing vascularity to improve healing of a segmental defect of the rat femur. Journal of orthopaedic trauma 2011, 25(8):472–476.

11. Zhang W, Li G, Deng R, Deng L, Qiu S: New bone formation in a true bone ceramic scaffold loaded with desferrioxamine in the treatment of segmental bone defect: a preliminary study. Journal of orthopaedic science: official journal of the Japanese Orthopaedic Association 2012, 17(3):289–298.

12. Wan C, Gilbert SR, Wang Y, Cao X, Shen X, Ramaswamy G, Jacobsen KA, Alaql ZS, Eberhardt AW, Gerstenfeld LC, Einhorn TA, Deng L, Clemens TL: Activation of the hypoxia-inducible factor-1alpha pathway accelerates bone regeneration. Proc Natl Acad Sci U S A 2008, 105(2):686–691.

13. Liu X, Tu Y, Zhang L, Qi J, Ma T, Deng L: Prolyl hydroxylase inhibitors protect from the bone loss in ovariectomy rats by increasing bone vascularity. Cell biochemistry and biophysics 2014, 69(1):141–149.

14. Peng J, Lai ZG, Fang ZL, Xing S, Hui K, Hao C, Jin Q, Qi Z, Shen WJ, Dong QN, Bing ZH, Fu DL: Dimethyloxalylglycine prevents bone loss in ovariectomized C57BL/6J mice through enhanced angiogenesis and osteogenesis. PloS one 2014, 9(11):e112744.

15. Joharapurkar AA, Pandya VB, Patel VJ, Desai RC, Jain MR: Prolyl Hydroxylase Inhibitors: A Breakthrough in the Therapy of Anemia Associated with Chronic Diseases. Journal of medicinal chemistry 2018.

16. Dhillon S: Roxadustat: First Global Approval. Drugs 2019, 79(5):563–572.

17. Fujikawa Y, Sabokbar A, Neale S, Athanasou NA: Human osteoclast formation and bone resorption by monocytes and synovial macrophages in rheumatoid arthritis. Ann Rheum Dis 1996, 55(11):816–822.

18. Quinn JM, Elliott J, Gillespie MT, Martin TJ: A combination of osteoclast differentiation factor and macrophage-colony stimulating factor is sufficient for both human and mouse osteoclast formation in vitro. Endocrinology 1998, 139(10):4424–4427.

19. Hulley PA, Bishop T, Vernet A, Schneider JE, Edwards JR, Athanasou NA, Knowles HJ: Hypoxia-inducible factor 1-alpha does not regulate osteoclastogenesis but enhances bone resorption activity via prolyl-4-hydroxylase 2. The Journal of pathology 2017, 242(3):322–333.

20. Knowles HJ, Athanasou NA: Hypoxia-inducible factor is expressed in giant cell tumour of bone and mediates paracrine effects of hypoxia on monocyte-osteoclast differentiation via induction of VEGF. J Pathol 2008, 215(1):56–66.

21. Knowles HJ, Athanasou NA: Acute hypoxia and osteoclast activity: a balance between enhanced resorption and increased apoptosis. J Pathol 2009, 218(2):256–264.

22. Knowles HJ, Cleton-Jansen AM, Korsching E, Athanasou NA: Hypoxia-inducible factor regulates osteoclast-mediated bone resorption: role of angiopoietin-like 4. The FASEB journal: official publication of the Federation of American Societies for Experimental Biology 2010, 24(12):4648–4659.

23. Miyauchi Y, Sato Y, Kobayashi T, Yoshida S, Mori T, Kanagawa H, Katsuyama E, Fujie A, Hao W, Miyamoto K, Tando T, Morioka H, Matsumoto M, Chambon P, Johnson RS, Kato S, Toyama Y, Miyamoto T: HIF1alpha is required for osteoclast activation by estrogen deficiency in postmenopausal osteoporosis. Proceedings of the National Academy of Sciences of the United States of America 2013, 110(41):16568–16573.

24. Rauner M, Franke K, Murray M, Singh RP, Hiram-Bab S, Platzbecker U, Gassmann M, Socolovsky M, Neumann D, Gabet Y, Chavakis T, Hofbauer LC, Wielockx B: Increased EPO Levels Are Associated With Bone Loss in Mice Lacking PHD2 in EPO-Producing Cells. Journal of bone and mineral research: the official journal of the American Society for Bone and Mineral Research 2016, 31(10):1877–1887.

25. Costa-Rodrigues J, Fernandes A, Fernandes MH: Reciprocal osteoblastic and osteoclastic modulation in co-cultured MG63 osteosarcoma cells and human osteoclast precursors. Journal of cellular biochemistry 2011, 112(12):3704–3713.

26. Heinemann S, Heinemann C, Wenisch S, Alt V, Worch H, Hanke T: Calcium phosphate phases integrated in silica/collagen nanocomposite xerogels enhance the bioactivity and ultimately manipulate the osteoblast/osteoclast ratio in a human co-culture model. Acta biomaterialia 2013, 9(1):4878–4888.

27. Chan MC, Ilott NE, Schodel J, Sims D, Tumber A, Lippl K, Mole DR, Pugh CW, Ratcliffe PJ, Ponting CP, Schofield CJ: Tuning the Transcriptional Response to Hypoxia by Inhibiting Hypoxia-inducible Factor (HIF) Prolyl and Asparaginyl Hydroxylases. The Journal of biological chemistry 2016, 291(39):20661–20673.

28. Zhou M, Hou J, Li Y, Mou S, Wang Z, Horch RE, Sun J, Yuan Q: The pro-angiogenic role of hypoxia inducible factor stabilizer FG-4592 and its application in an in vivo tissue engineering chamber model. Scientific reports 2019, 9(1):6035.

29. Li X, Cui XX, Chen YJ, Wu TT, Xu H, Yin H, Wu YC: Therapeutic Potential of a Prolyl Hydroxylase Inhibitor FG-4592 for Parkinson’s Diseases in Vitro and in Vivo: Regulation of Redox Biology and Mitochondrial Function. Frontiers in aging neuroscience 2018, 10:121.

30. Xie RY, Fang XL, Zheng XB, Lv WZ, Li YJ, Ibrahim Rage H, He QL, Zhu WP, Cui TX: Salidroside and FG-4592 ameliorate high glucose-induced glomerular endothelial cells injury via HIF upregulation. Biomedicine & pharmacotherapy = Biomedecine & pharmacotherapie 2019, 118:109175.

31. Mokas S, Lariviere R, Lamalice L, Gobeil S, Cornfield DN, Agharazii M, Richard DE: Hypoxia-inducible factor-1 plays a role in phosphate-induced vascular smooth muscle cell calcification. Kidney international 2016, 90(3):598–609.

32. Matthews BG, Naot D, Callon KE, Musson DS, Locklin R, Hulley PA, Grey A, Cornish J: Enhanced osteoblastogenesis in three-dimensional collagen gels. BoneKEy reports 2014, 3:560.

33. Gurski LA, Jha AK, Zhang C, Jia X, Farach-Carson MC: Hyaluronic acid-based hydrogels as 3D matrices for in vitro evaluation of chemotherapeutic drugs using poorly adherent prostate cancer cells. Biomaterials 2009, 30(30):6076–6085.

34. Sawa N, Fujimoto H, Sawa Y, Yamashita J: Alternating Differentiation and Dedifferentiation between Mature Osteoblasts and Osteocytes. Scientific reports 2019, 9(1):13842.

35. Park HJ, Baek KH, Lee HL, Kwon A, Hwang HR, Qadir AS, Woo KM, Ryoo HM, Baek JH: Hypoxia inducible factor-1alpha directly induces the expression of receptor activator of nuclear factor-kappaB ligand in periodontal ligament fibroblasts. Molecules and cells 2011, 31(6):573–578.

36. Tang ZN, Zhang F, Tang P, Qi XW, Jiang J: Hypoxia induces RANK and RANKL expression by activating HIF-1alpha in breast cancer cells. Biochem Biophys Res Commun 2011, 408(3):411–416.

37. Zhu J, Tang Y, Wu Q, Ji YC, Feng ZF, Kang FW: HIF-1alpha facilitates osteocyte-mediated osteoclastogenesis by activating JAK2/STAT3 pathway in vitro. J Cell Physiol 2019, 234(11):21182–21192.

38. Lee SY, Park KH, Yu HG, Kook E, Song WH, Lee G, Koh JT, Shin HI, Choi JY, Huh YH, Ryu JH: Controlling hypoxia-inducible factor-2alpha is critical for maintaining bone homeostasis in mice. Bone research 2019, 7:14.

39. Madel MB, Ibanez L, Rouleau M, Wakkach A, Blin-Wakkach C: A Novel Reliable and Efficient Procedure for Purification of Mature Osteoclasts Allowing Functional Assays in Mouse Cells. Frontiers in immunology 2018, 9:2567.

40. Abdallah D, Jourdain ML, Braux J, Guillaume C, Gangloff SC, Jacquot J, Velard F: An Optimized Method to Generate Human Active Osteoclasts From Peripheral Blood Monocytes. Frontiers in immunology 2018, 9:632.

41. Wu L, Feyerabend F, Schilling AF, Willumeit-Romer R, Luthringer BJC: Effects of extracellular magnesium extract on the proliferation and differentiation of human osteoblasts and osteoclasts in coculture. Acta biomaterialia 2015, 27:294–304.

42. Kim JA, Ihn HJ, Park JY, Lim J, Hong JM, Kim SH, Kim SY, Shin HI, Park EK: Inhibitory effects of triptolide on titanium particle-induced osteolysis and receptor activator of nuclear factor-kappaB ligand-mediated osteoclast differentiation. International orthopaedics 2015, 39(1):173–182.

43. Costa-Rodrigues J, Carmo SC, Silva JC, Fernandes MH: Inhibition of human in vitro osteoclastogenesis by Equisetum arvense. Cell proliferation 2012, 45(6):566–576.

44. Nishide S, Matsunaga S, Shiota M, Yamaguchi T, Kitajima S, Maekawa Y, Takeda N, Tomura M, Uchida J, Miura K, Nakatani T, Tomita S: Controlling the Phenotype of Tumor-Infiltrating Macrophages via the PHD-HIF Axis Inhibits Tumor Growth in a Mouse Model. iScience 2019, 19:940–954.

45. Xia L, Kilb J, Wex H, Li Z, Lipyansky A, Breuil V, Stein L, Palmer JT, Dempster DW, Bromme D: Localization of rat cathepsin K in osteoclasts and resorption pits: inhibition of bone resorption and cathepsin K-activity by peptidyl vinyl sulfones. Biological chemistry 1999, 380(6):679–687.

46. Blair HC, Sidonio RF, Friedberg RC, Khan NN, Dong SS: Proteinase expression during differentiation of human osteoclasts in vitro. J Cell Biochem 2000, 78(4):627–637.

47. Bongio M, Lopa S, Gilardi M, Bersini S, Moretti M: A 3D vascularized bone remodeling model combining osteoblasts and osteoclasts in a CaP nanoparticle-enriched matrix. Nanomedicine 2016, 11(9):1073–1091.

